# Woolly Ash Aphid *Prociphilus Fraxinifolii* (Riley, 1879) (Hemiptera: Eriosomatidae), A New Invasive Alien Pest of Ash Trees (*Fraxinus*) in Russia

**DOI:** 10.1101/254078

**Authors:** Andrzej Bieńkowski, Marina Orlova-Bienkowskaja

## Abstract

Woolly ash aphid or ash leaf curl aphid *Prociphilus fraxinifolii* (Riley, 1879) is an alien invasive pest of ash trees native to North America. After its first record in Europe in 2003 in Hungary it has spread to the Ukraine, Serbia, Bulgaria, Great Britain, Spain, Poland and Germany. In 2016 *P. fraxinifolii* was firstly recorded at the southwestern border of Russia. Now *Prociphilus fraxinifolii* is firstly recorded in the center of European Russia, namely in Moscow Region, which is more than 700 km far from all other known localities of the species. In September 2017 five groups of ash trees (*Fraxinus pennsylvanica*) with colonies of *Prociphilus fraxinifolii* were found in Moscow Region. The example of *Prociphilus fraxinifolii* shows that alien pest insects can spread in Europe very quickly. Now Moscow region is the only regions of Europe, where the expanding range of *Prociphilus fraxinifolii* has overlapped with expanding ranges of other invasive alien species established in Europe in the last 20 years and severely damaging ash trees: ash dieback fungus *Hymenoscyphus fraxineus* Baral et al., 2014 (Ascomycota) and emerald ash borer *Agrilus planipennis* Fairmaire, 1888 (Coleoptera: Buprestidae).

## INTRODUCTION

Ash trees (*Fraxinus* spp.) comprise a significant part of urban plantations and forests in Europe and play an important role in biodiversity, since many animals, plants and fungi are connected with them (Fraxigen, 2005). In the last two decades ash trees in Europe are affected by new alien serious pests and pathogens: ash dieback fungus *Hymenoscyphus fraxineus* Baral et al., 2014 (Ascomycota), emerald ash borer *Agrilus planipennis* Fairmaire, 1888 (Coleoptera: Buprestidae) and ash leaf curl aphid *Prociphilus fraxinifolii* (Riley, 1879) (Hemiptera: Eriosomatidae) (Herms & McCullough, 2014; Gross *et al*., 2014; Hałaj & Osiadacz, 2017). They are spreading quickly and causing significant damage to the trees. All these harmful species were found in Europe less than 20 years ago, all are spreading quickly and causing significant damage to the trees. Here we report the new data on further spread of *P. fraxinifolii* to the east.

Woolly ash aphid or ash leaf curl aphid *Prociphilus fraxinifolii* is native to North America. It is a common pest of ash trees in the United States, Canada and Mexico and is introduced to Chile, South Africa, China, Iran and Europe (Hałaj & Osiadacz, 2017). In 2003 it was firstly recorded in Europe in Budapest (Hungary) (Remaudiere & Ripka, 2003). Then it quickly spread to the Ukraine (first record in 2005), Serbia (2006), Bulgaria (2007), Great Britain (2011), Spain (2011), Poland (2012), and Germany (2015) (Martynov & Nikulina, 2016; Hałaj & Osiadacz, 2017). In 2016 *P. fraxinifolii* was firstly recorded at the southwestern border of Russia (Rostov Region, Avilo-Uspenka Village) (Martynov & Nikulina, 2016). Here *P. fraxinifolii* is firstly recorded in the center of European Russia, namely in Moscow Region, which is more than 700 km far from all other known localities of the species.

## SURVEY OF ASH TREES AND COLLECTION OF MATERIAL

*Fraxinus pennsylvanica* introduced from North America is widely cultivated in European Russia since the 1950s. Now it makes up about 20% of the trees in urban plantations of Central Russia, in particular, in Moscow (Majorov *et al*., 2012). The outbreak of the emerald ash borer *Agrilus planipennis* on *F. pennsylvanica* prompted us to study the community of insects connected with this tree in European Russia. In 2013–2017 we surveyed more than 6000 ash trees in 40 localities in 20 regions of European Russia (Orlova-Bienkowskaja, 2014, 2015; Orlova-Bienkowskaja & Volkovitsh, 2014; Orlova-Bienkowskaja & Belokobylskij, 2014; Orlova-Bienkowskaja & Bieńkowski, 2016 and unpublished data). Besides this, regular surveys of ash trees were carried out in Moscow and Zelenograd (Moscow Region) in 2013–2017. Characteristic pseudogalls with colonies *P. fraxinifolii* were firstly detected during survey of *F. pennsylvanica* trees on 11 September 2017 in two localities in Moscow: near Mitino metro station (three trees) and at the intersection of Moscow Ring Road and Volokolamskoe Highway (two trees). After that in September 2017 a special survey of ash trees was carried out in Zelenograd, where *F. pennsylvanica* is very common in all districts. About 1000 trees of *F. pennsylvanica* have been examined. Three groups of trees damaged by these aphids have been found in different districts of the city: two trees in the 11th district, three trees in the 4th district and two trees and the 2nd district. Groups of damaged trees occurred in the same plantations along the streets with the trees unaffected by the pest.

All trees, where aphids were found, have emergence holes of *Agrilus planipennis,* like the most of *F. pennsylvanica* trees in Moscow Region. But the canopies of these trees were not thinned. Up to 20 pseudogalls were observed on each tree. Pseudogall is a deformed shortened shoot with tightly curled leaves, which is a “nest” for aphid colony consisting of apterae and alatae viviparae (Fig. I). A colony is immersed in the thick wax produced by insects. Aphids excrete large amounts of white honeydew, which is seen on the leaves under the “nests”. Collected aphids were placed to the alcohol. This material is deposited in the collection of the first author (Zelenograd). Slides of alatae and apterous females were prepared and examined under the microscope.

**Fig. I.**
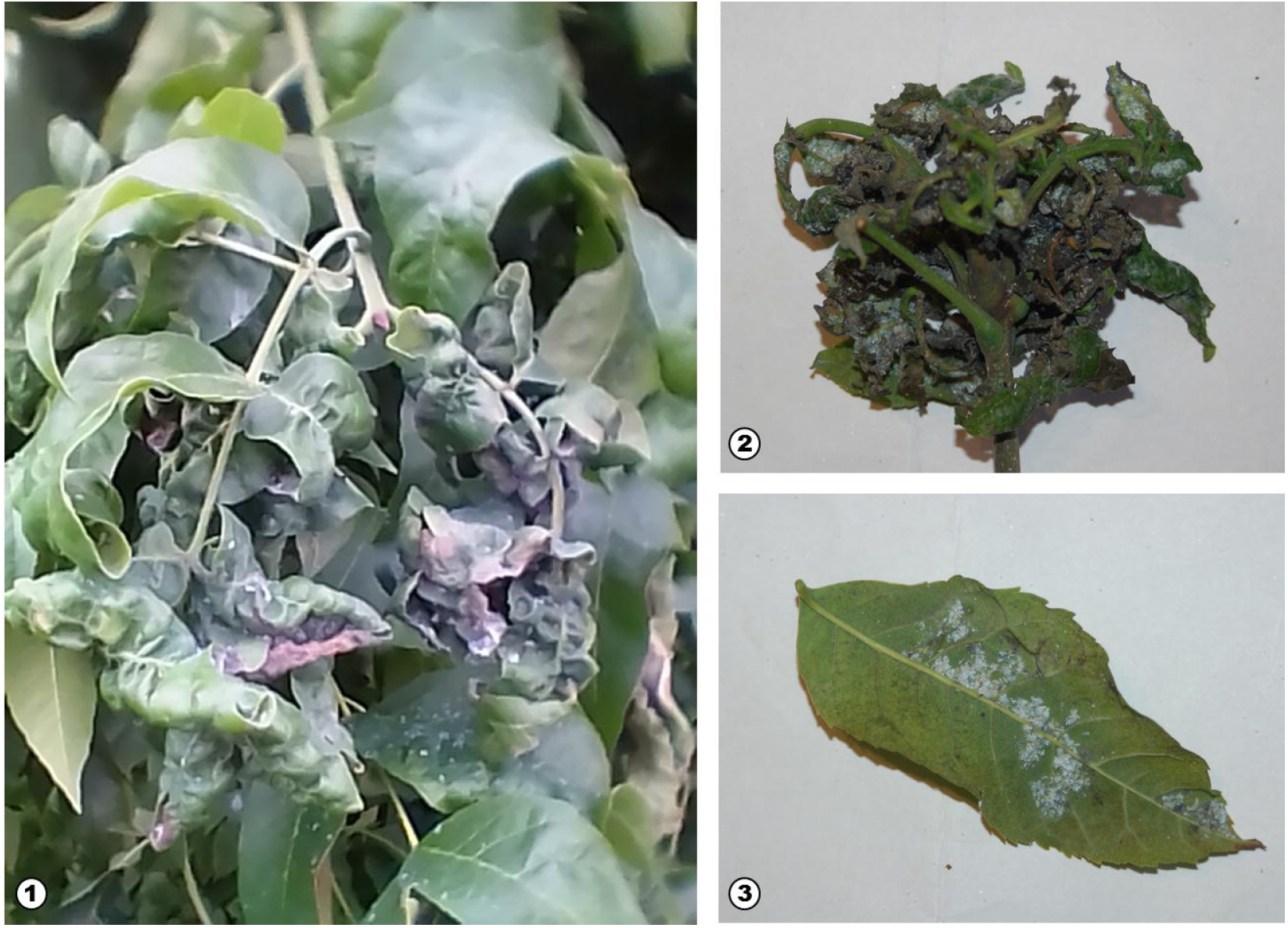
*Prociphilus fraxinifolii* on *Fraxinus pennsylvanica* in Moscow Region (Russia). 1,2 – Malformations or pseudogalls caused by these aphids; 3 – leaf with colony of *P. fraxinifolii.*

## IDENTIFICATION

The species was identified referring to Blackman and Eastop (2016). The genus *Prociphilus* Koch, 1856 includes about 45 species grouped into six subgenera (Blackman & Eastop, 2016). Subgenus *Prociphilus* (*Meliarhizophagus*) Smith, 1974 consists of only one species – *Prociphilus* (*Meliarhizophagus*) *fraxinifolii.*

Details of adult apterous female and alatae female are shown in Fig. II. *Prociphilus* (*Meliarhizophagus*) *fraxinifolii* can be distinguished from other species of the genus by the following characters. Alatae females have secondary rhinaria on antennomere 6 differing in shape from narrow transverse secondary rhinaria on antennomere 3. Antennomere 3 as long as 3x antennomere 2. Besides that, they differ from those of *P. bumeliae*, and *P. fraxini* in antennomere 5 with 4 secondary rhinaria, antennomere 6 with 2 irregular-rounded secondary rhinaria (1-5 rhinaria after Blackman & Eastop, 2016). Apterous mature females of *P. fraxinifolii* differ from morphologically close *P. probosceus* in rostrum very short, with apical segment (really 4^th^ + 5^th^ segments joined together) with 3 accessory hairs (2 hairs after Blackman & Eastop, 2016).

**Fig. II.**
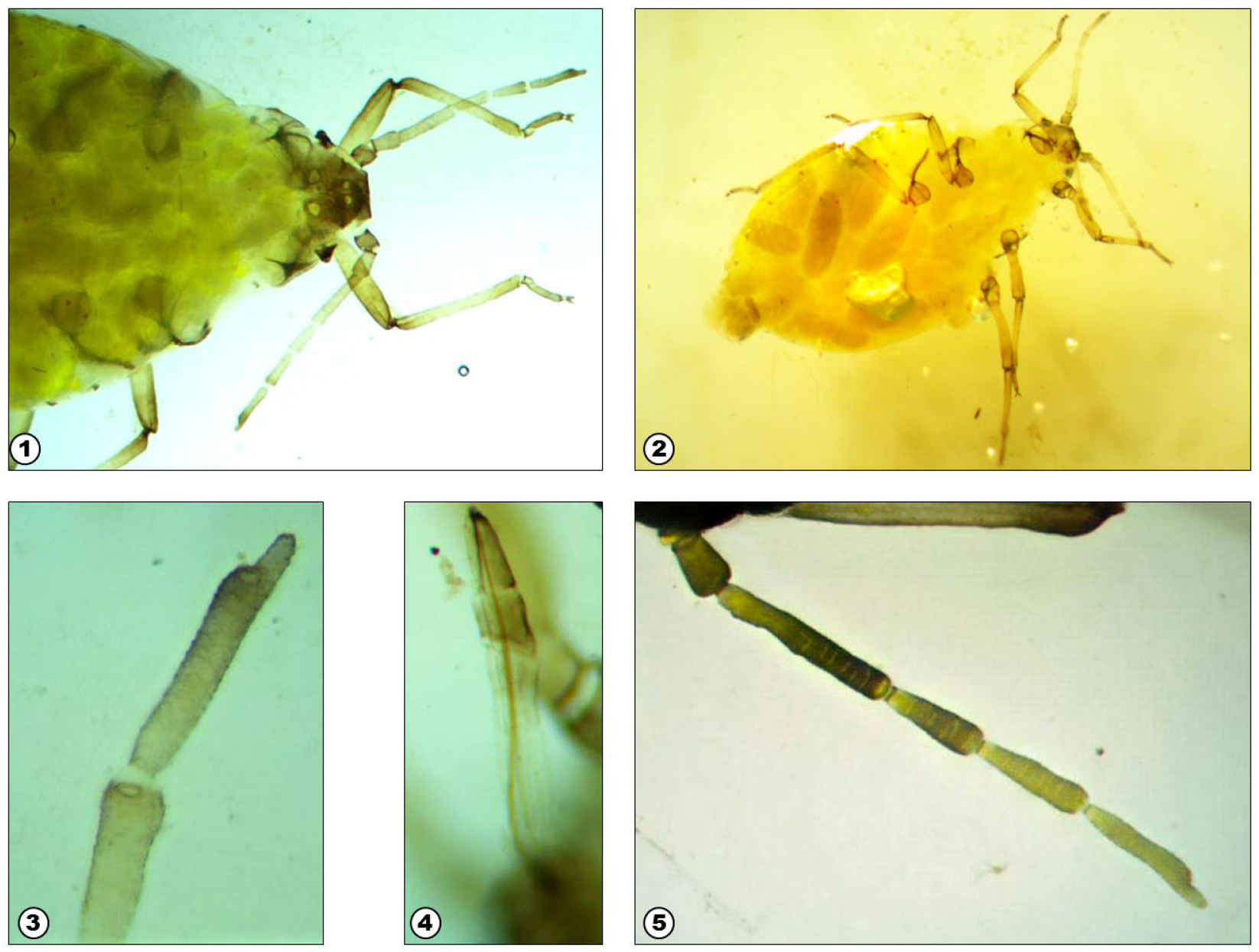
*Prociphilus fraxinifolii* on *Fraxinus pennsylvanica* in Moscow Region (Russia). 1 – anterior part of the body of adult apterous female from above, 2 – apterous female with embryons from below, 3 – last antennal segment of apterous female, 4 – rostrum of apterous female, antenna of alatae female.

Mature and immature apterous viviparous females of *P. fraxinifolii* are pale green or pale yellow. Nymphs pale green with thorax pale yellow. Alatae viviparous females have grey head and thorax and pale green abdomen. Ocelli and compound eyes (in alatae females and nymphs) are black. All morphs are covered by white wax, besides that, mature and immature apterous females bear thick curved white wax threads.

## BIOLOGY AND HARMFULNESS

In its native range *P. fraxinifolii* feeds on several North American species of the genus *Fraxinus*. Outside the native range *P. fraxinifolii* was initially recorded only on species introduced from North America: *F. pennsylvanica, F. americana*, and *F. velutina*, so it was suggested that *P. fraxinifolii* did not affect native ash trees in Europe (Coeur d’acier *et al*., 2010; Pérez Hidalgo & Mier Durante, 2012). But then *P. fraxinifolii* was also found on European ash *F. excelsior* (Hałaj & Osiadacz, 2017). We have yet found *P. fraxinifolii* only on *F. pennsylvanica,* because *F. excelsior* is rare in Moscow.

The genus *Prociphilus* is connected mainly with the trees of the genus *Fraxinus*. Four species besides *P. fraxinifolii* are recorded on *Fraxinus pennsylvanica* in the world: *P.* (*Prociphilus*) *americanus* (Walker, 1852), *Prociphilus* (*Pulvius*) *probosceus* Sanborn, 1904*, Prociphilus* (*Prociphilus*) *bumeliae* (Schrank, 1801) and *Prociphilus* (*Prociphilus*) *fraxini* (Fabricius, 1777). *P. americanus* and *P. probosceus* occur only in America. *Prociphilus bumeliae* and *P. fraxini* are native European species connected mainly with native European ash *Fraxinus excelsior* and sometimes occurring also on the introduced ash *F. pennsylvanica* (Blackman & Eastop, 2016).

*Prociphilus fraxinifolii* is monoecious, i.e. does not change host plants during its life cycle (Hałaj & Osiadacz, 2017). This feature distinguishes it from other European representatives of the genus feeding on ash trees. *Prociphilus fraxini* and *P. bumeliae* migrate from *Fraxinus* to *Abies* in summer (Hałaj *et al*., 2016).

The opinions about the harmfulness of *P. fraxinifolii* are contradicting. Pérez Hidalgo and Mier Durante (2012) suppose that these aphids do not seem to cause any significant harm to trees. But Hałaj and Osiadacz (2017) argue that *P. fraxinifolii* can significantly damage the tree. Our observations confirm the latter point of view. First, the formation of pseudogalls reduces the growth of shoots and causes their premature dieback. This has a significant negative impact, since mainly young trees are affected and up to 20 pseudogalls appear on one tree. Second, the large amount of honeydew produced by *P. fraxinifolii* facilitates the development of harmful fungi. In about all of the trees affected by these aphids in Moscow Region the leaves were also strongly affected by powdery mildew. The appearance of *P. fraxinifolii* is especially dangerous to the trees which are already weakened by the emerald ash borer.

Predators, in particular, ladybirds, easily penetrate to the pseudogalls. It is known that the outbreak of *P. fraxinifolii* sometimes facilitates the outbreak of ladybirds (Torres-Acosta & Sánchez-Peña, 2015). The following coccinellids have been found on *F. pannsylvanica* affected by *P. fraxinifolii* in Moscow Region: *Adalia bipunctata* (*Linnaeus, 1758*), *Psyllobora vigintiduopunctata* (Linnaeus, 1758), *Exochomus quadripustulatus* (Linnaeus, 1758), *Calvia decemguttata* (Linnaeus, 1767) and *Halyzia sedecimguttata* (Linnaeus, 1758).

## DISCUSSION

*Prociphilus fraxinifolii* is spreading in Europe very quickly. The spread of *P. fraxinifolii* in the eastern part of the range has been described by Martynov and Nikulina (2016). In 2005, in only two years after the first finding in Europe in Budapest, the pest was recorded in the west of the Ukraine – in Uzhgorod (Transcarpathian region). In 2012 it appeared in the center of the Ukraine: in Kiev. Then in 2015 it was firstly recorded in the east of the country: in Donetsk and Lugansk Regions. In summer 2016 the pest became very common in urban plantations of Donetsk and was firstly recorded at the southeastern border of Russia in Rostov Region. Now *P. fraxinifolii* is recorded much further north. The distance between Moscow and the nearest localities of *P. fraxinifolii* known before is more than 700 km. It indicates that the dispersal of the pest to the center of European Russia was not natural, but represented an accidental introduction by human. It is not surprising that the pest was introduced just to Moscow. It is the largest city of Europe, the main transport hub of Russia, and therefore the main center of invasions of insects. Alien insects are often introduced to this city and after the establishment quickly spread to other regions (Orlova-Bienkowskaja, 2017).

It seems that *P. fraxinifolii* appeared in the center of European Russia just recently, because it was not found in 2013–2016 in spite of our intensive surveys of ash trees in different regions, especially in Moscow Region. The characteristic pseudogalls are easy to notice, but they were not found until the autumn 2017. Now the pest has not yet become very abundant in Moscow Region. Only individual foci (small groups of trees affected by the pest) are found in Moscow Region, and only about 1% of surveyed trees are affected by the pest. But it is quite possible that the strong outbreak will happen in the near future as it was observed in Donetsk (Martynov & Nikulina, 2016). The first individual foci of the pest were found in Donetsk in 2015. But in summer 2016 *P. fraxinifolii* became very common and abundant on ash trees in all types of plantations all over the city (Martynov & Nikulina, 2016).

It is interesting that three alien major pests and pathogens of ash trees appeared in Europe at the same time. Ash dieback fungus *Hymenoscyphus fraxineus* appeared in Poland in the end of 1990s (Kowalski, 2006), emerald ash borer *Agrilus planipennis* was recorded in Moscow in 2003 (Shankhiza, 2007), and *P. fraxinifolii* was recorded in Hungary in 2003 (Remaudiere & Ripka, 2003). These species could not be introduced together, since the first and second ones are native to East Asia, though the third one originates from North America. But it is hard to believe that almost simultaneous establishment of three alien species severely damaging ash trees is just a coincidence. It is more likely that it was facilitated by weakening the trees by some factor, for example, by climatic changes. It is also probably that these three species “helped” each other to become established by weakening the trees. Such positive interactions of nonnative species are known as an invasional meltdown (Simberloff & Von Holle, 1999). Special plant protection and plant quarantine measures should be taken to minimize negative impact of established alien species connected with ash trees in Europe and to prevent establishment of new alien pest and pathogens of these trees.

By now Moscow Region has become the only region where all three alien species seriously damaging ash trees are recorded: *Agrilus planipennis* appeared in 2003, *Hymenoscyphus fraxineus* reached Moscow by 2014 (Musolin *et al*., 2017) and *P. fraxinifolii* reached Moscow by 2017 (current communication).

## ACKNOWLEDGMENTS

The study was supported by Russian Science Foundation, Project No 16-14-10031.

## REFERENCES

Coeur D’Acier A., Pérez Hidalgo N., Petrović-Obradović O., 2010 – *Aphids (Hemiptera, Aphididae).* Chapter 9.2. A. Roques, M. Kenis, D. Lees, C. Lopez-Vaamonde, W. Rabitsch, J.-Y. Rasplus & D.B. Roy Ed., BioRisk. 4 (1). Pensoft, Sofia – Moscow pp. 435–474.

Blackman R.L., Eastop V.F., 2016 - Aphids on the world’s plants. An online identification and information guide. URL:http://www.aphidsonworldsplants.info [accessed Sept 17, 2017].

FRAXIGEN, 2005 – Ash species in Europe: biological characteristics and practical guidelines for sustainable use. Oxford Forestry Institute, University of Oxford, UK, 128 pp. URL: http://herbaria.plants.ox.ac.uk/fraxigen/pdfs_and_docs/book/fraxigen_c1toc3.pdf.

Gross A., Holdenrieder O., Pautasso M., Queloz V., Sieber T.N., 2014 – Hymenoscyphus pseudoalbidus, *the causal agent of European ash dieback*. - Molecular Plant Pathology, 15 (1): 5–21, DOI: 10.1111/mpp.12073.

Hałaj R., Osiadacz B., Poljaković-Pajnik L., 2016 – *Górny Śląsk-polski przyczółek zdobyty przez obcą mszycę* Prociphilus (Meliarhizophagus) fraxinifolii (Riley, 1879). - Acta Entomologica Silesiana, 24: 1–9.

Hałaj R., Osiadacz B., 2017 – Woolly ash aphid – is the alien bug posing a threat to European ash trees? – a review. - Plant Protection Science 53 (3): 127–133, DOI: 10.17221/138/2016-PPS.

Herms D.A., McCullough D.G., 2014 – Emerald ash borer invasion of North America: history, biology, ecology, impacts, and management. - Annual Review of Entomology 59: 13–30, DOI: 10.1146/annurev-ento-011613-162051

Kowalski T., 2006 – Chalara fraxinea *sp. nov. associated with dieback of ash* (Fraxinus excelsior) *in Poland*. - Forest Pathology, 36 (4): 264–270.

Majorov S.R., Bochkin V.D., Nasimovich Y.A., Shcherbakov A.V., 2012 – Alien Flora of Moscow and Moscow Region. KMK, Moscow, 412 pp.

Martynov V., Nikulina T., 2016 – Prociphilus (Meliarhizophagus) *fraxinifolii (Riley, 1979) (Hemiptera: Aphididae: Eriosomatinae) – a new invasive North American species of aphids in the territory of Donbass*, Actual problems of integrated plant protection, Kyiv, pp. 53–54.

Musolin D.L., Selikhovkin A.V., Shabunin D.A., Zviaginsev V.B., Baranchikov Y.N., 2017 – Between ash dieback and emerald ash borer: two Asian invaders in Russia and the future of ash in Europe. - Baltic Forestry, 23 (1): 316–333.

Orlova-Bienkowskaja M.J., 2014 – *Ashes in Europe are in danger: the invasive range of* Agrilus planipennis *in European Russia is expanding*. - Biological Invasions, 16: 1345–1349. DOI: 10.1007/s10530-013-0579-8

Orlova-Bienkowskaja M.J., 2015 – *Cascading ecological effects caused by establishment of the emerald ash borer* Agrilus planipennis *in European Russia*. - European Journal of Entomology 112 (4): 778–789, DOI: 10.14411/eje.2015.102

Orlova-Bienkowskaja M.J., 2017 – Main Trends of Invasion Processes in Beetles (Coleoptera) of European Russia. - Russian Journal of Biological Invasions, 8 (2): 143–157, DOI: 10.1134/S2075111717020060

Orlova-Bienkowskaja M.J., Belokobylskij S.A., 2014 – *Discovery of the first European parasitoid of the emerald ash borer* Agrilus planipennis *Fairmaire (Coleoptera: Buprestidae)*. - European Journal of Entomology, 111 (4): 594–596, DOI: 10.14411/eje.2014.061

Orlova-Bienkowskaja M.J., Bieńkowski A.O., 2016 – *The life cycle of the emerald ash borer* Agrilus planipennis *in European Russia and comparisons with its life cycles in Asia and North America*. - Agricultural and Forest Entomology, 18 (2): 182–188, DOI: 10.1111/afe.12140

Orlova-Bienkowskaja M.J., Volkovitsh M.G., 2014 – *Range expansion of* Agrilus convexicollis *in European Russia expedited by the invasion of emerald ash borer,* Agrilus planipennis (Coleoptera: Buprestidae). - Biological Invasions, 17 (2): 537–544, DOI: 10.1007/s10530-014-0762-6

Pérez Hidalgo N., Mier Durante M.P., 2012 – *First record of* Prociphilus (Meliarhizophagus) fraxinifolii *(Riley) [Hemiptera: Aphididae] in the Iberian Peninsula*. - EPPO Bulletin, 42 (1): 142–145, DOI: 10.1111/epp.2531

Remaudiere G., Ripka G., 2003 – *Arrivée en Europe (Budapest, Hongrie) du puceron des frenes américains,* Prociphilus (Meliarhizophagus) fraxinifolii *(Hemiptera, Aphididae, Eriosomatinae, Pemphigini)*. - Revue française d’entomologie, 25 (3): 152.

Shankhiza E.V., 2007 - Invasion of the emerald ash borer Agrilus planipennis to Moscow Region. – URL: http://www.zin.ru/Animalia/Coleoptera/rus/fraxxx.htm [accessed Sept 17, 2017].

Simberloff D., Von Holle B., 1999 – Positive interactions of nonindigenous species: invasional meltdown? - Biological Invasions.1: 21–32.

Torres-Acosta R. I., Sánchez-Peña, S. R., 2015 – *Regional concurrent outbreaks of ash leaf curl aphid,* Prociphilus fraxinifolii *(Riley) (Hemiptera: Aphididae: Eriosomatinae), and the invasive predator,* Harmonia axyridis *(Pallas) (Coleoptera: Cocinellidae), in Northeastern Mexico*. - Southwestern Entomologist, 40 (3): 661–664.

